# Amyloid-β, p-tau, and reactive microglia load are correlates of MRI cortical atrophy in Alzheimer’s disease

**DOI:** 10.1101/2021.06.16.448650

**Authors:** Irene Frigerio, Baayla DC Boon, Chen-Pei Lin, Yvon Galis-de Graaf, John GJM Bol, Paolo Preziosa, Jos Twisk, Frederik Barkhof, Jeroen JM Hoozemans, Femke H Bouwman, Annemieke JM Rozemuller, Wilma DJ van de Berg, Laura E Jonkman

## Abstract

**INTRODUCTION:** The aim of this study was to identify the histopathological correlates of MRI cortical atrophy in (a)typical Alzheimer’s disease (AD) donors.

**METHODS:** 19 AD and 10 control donors underwent post-mortem *in-situ* 3T-3DT1-MRI, from which cortical thickness was calculated. Upon subsequent autopsy, 21 cortical brain regions were selected and immunostained for amyloid-beta, phosphorylated-tau, and reactive microglia. MRI-pathology associations were assessed using linear mixed models. Post-mortem MRI was compared to ante-mortem MRI when available.

**RESULTS:** Higher amyloid-beta load weakly correlated with a higher cortical thickness globally. Phosphorylated-tau strongly correlated with cortical atrophy in temporo-frontal regions. Reactive microglia load strongly correlated with cortical atrophy in the parietal region. Post-mortem scans showed high concordance with ante-mortem scans acquired <1 year before death.

**DISCUSSION:** Distinct histopathological markers differently correlate with cortical atrophy, highlighting their different roles in the neurodegenerative process. This study contributes in understanding the pathological underpinnings of MRI atrophy patterns.

## 1. BACKGROUND

Alzheimer’s disease (AD) is a progressive neurodegenerative disease with a heterogeneous clinical presentation. Clinically, AD is defined as typical when memory deficits are the first complaints, and atypical when memory is initially spared while other symptoms are more prominent, such as visuospatial impairment, aphasia, or behavioural/dysexecutive dysfunction^1^. On magnetic resonance imaging (MRI), typical AD is typically characterized by hippocampal and temporoparietal atrophy^2^, whereas atypical presentations may show initial hippocampal sparing, and cortical atrophy in regions corresponding to clinical symptoms^3^. Although hippocampal and temporoparietal atrophy are commonly used MRI biomarkers in AD, reliable imaging biomarkers ideally reflect disease state as well as underlying pathophysiological mechanisms in AD, and should be validated in a cohort of neuropathologically-confirmed AD subjects.

Pathologically, AD is characterised by the accumulation of amyloid-beta (Aβ) plaques and phosphorylated-tau (p-tau) neurofibrillary tangles (NFT) in the grey matter^4,5^. In addition, neuroinflammation plays a key role in AD^6,7^. Reactive microglia tend to proliferate and increase with disease progression^6,7^ and seem to reflect p-tau deposition sites^8^.

Studies combining MRI and positron emission tomography (PET) imaging as a proxy for pathology showed that the load and neuroanatomical distribution of tau tracers correlate with cortical atrophic patterns on MRI^9–13^, while widespread cortical Aβ deposition^9^ does not correlate with cortical atrophy nor clinical presentation in clinically defined AD cases^14,15^. The effect of reactive microglia load on cortical atrophy remains elusive, since the few studies that investigated its association with MRI atrophic patterns report contrasting results^16,17^. Unfortunately, PET imaging cannot evaluate different pathological processes together, but just one at each exam. As such, post-mortem histopathological examination remains the gold-standard to diagnose AD and to evaluate the combination of different markers^18^.

The aim of the current study was to assess the association between post-mortem *in-situ* (within the skull) MRI cortical thickness and histopathological hallmarks in clinically-defined and pathologically-confirmed AD subjects. Additionally, due to the clinical heterogeneity in AD, we explored the association between neuroanatomical distribution of MRI and pathological patterns in clinically-defined typical and atypical AD phenotypes^19^. Lastly, we investigated the coherence between ante-mortem and post-mortem patterns in a subsets of these patients. The results of this study will increase our knowledge on the histopathological correlates of cortical atrophy, thereby contributing to the understanding of the pathological underpinnings of MRI atrophy patterns.

## 2. METHODS

Detailed methods are described in Supplementary Methods.

### 2.1 Donor inclusion

In collaboration with the Netherlands Brain Bank (NBB; http://brainbank.nl) we included 19 AD donors from the Amsterdam Dementia Cohort^20^. The AD donors could be further subdivided into 10 typical and 9 atypical AD donors based on clinical symptoms, of which 6 were diagnosed with the behavioural/dysexecutive variant (B/D)^3^, and 3 with posterior cortical atrophy (PCA)^21^. Neuropathological diagnosis was confirmed by an expert neuropathologist (A.R.) and performed according to the international guidelines of the Brain Net Europe II (BNE) consortium (http://www.brainnet-europe.org)^22,23^. Additionally, 10 age-matched pathologically confirmed non-neurological controls were selected from the Normal Aging Brain Collection Amsterdam (NABCA; http://nabca.eu)^24^. All donors signed an informed consent for brain donation and the use of material and clinical information for research purposes. The procedures for brain tissue collection of NBB and NABCA have been approved by the Medical Ethical Committee of VUmc. For donor characteristics, see **Table S1**.

### 2.2 Post-mortem *in-situ* and ante-mortem *in-vivo* MRI acquisition

Post-mortem 3T brain *in-situ* MRI acquisition was acquired according to a previously described pipeline^25^ and explained in detail in the Supplementary Methods. Briefly, 3T MRI was acquired on a magnetic resonance scanner (Signa-MR750, General Electric Medical Systems, United States) with an eight-channel phased-array head-coil. The cortical surface was reconstructed with FreeSurfer, version 6.0 (http://surfer.nmr.mgh.harvard.edu)^26^. Moreover, 14 AD out of 19 patients included in our study had ante-mortem *in-vivo* 3T MRI scans available, and these were included for comparison with post-mortem MRI. Details about these scans can be found in Table S2.

### 2.3 Tissue sampling

Formalin-fixed paraffin-embedded (4%, 4 weeks fixation) tissue blocks from the following regions of the right hemisphere were used: superior and middle frontal gyrus, anterior and posterior cingulate gyrus, middle temporal gyrus, superior and inferior parietal gyrus, precuneus, occipital cortex (primary visual cortex), and hippocampus (including the entorhinal cortex, parahippocampal and fusiform gyrus as described before^27^). Additionally, for 13 AD cases (7 typical and 6 atypical), 9 formalin-fixed paraffin-embedded (4%, 24-36 hours fixation) tissue blocks from the left hemisphere were available from the same regions as described above except for the hippocampus and posterior cingulate gyrus, adding the superior temporal gyrus.

### 2.4 Immunohistochemistry and image analysis

The immunohistological procedures are explained in detail in the Supplementary Methods. Briefly, 6-µm thick sections were cut from the above-mentioned regions. The sections from the right hemisphere were stained for Aβ (4G8), p-tau (AT8), and reactive microglia (CD68, clone KP1). The sections from the left hemisphere were additionally collected and stained for Aβ (4G8) and p-tau (AT8) (see Table S3 for information on primary antibodies). Images were taken using a whole-slide scanner (Vectra Polaris, 20x objective) and quantified using Fiji ImageJ Version 1.52r (https://imagej.nih.gov/ij). Regions of interest (ROIs) containing all cortical layers were delineated in straight areas of the cortex to avoid over- or underestimation of pathology in sulci and gyri, respectively^28^. After colour deconvolution, used to separate haematoxylin and 3.3’-Diaminobenzidine (DAB) channels, the immunoreactivity of DAB staining was quantified using the auto-threshold plugin “maximum entropy”^29^. The outcome measure was the %DAB-stained area per ROIs of each section per marker. For an overview of our workflow, see Figure 1.

**Figure 1.**
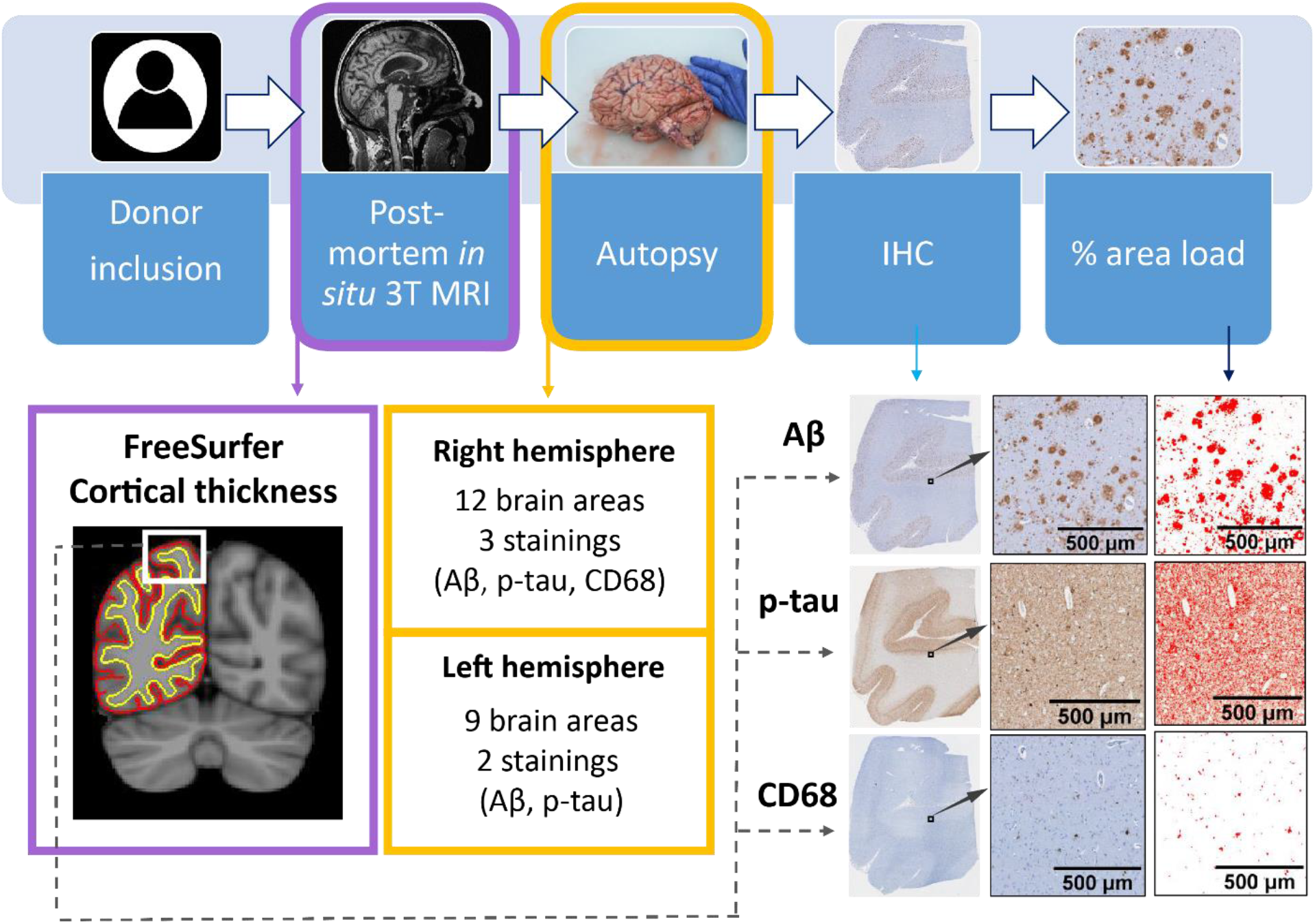
Workflow of the post-mortem MRI-pathology pipeline. Once the donors were included in the study, they received a post-mortem *in-situ* 3T MRI, and cortical thickness was calculated with FreeSurfer^26^ from the 3D T1w image (purple box). After the MRI scan, autopsy was performed, and brain tissue was processed for immunohistochemistry against Aβ, p-tau and CD68 (yellow boxes), which were quantified using ImageJ. The correlation between cortical thickness and %area of immunoreactivity was investigated via linear mixed models (dashed grey arrow). Legend: IHC = immunohistochemistry; Aβ = amyloid-beta; p-tau = phosphorylated-tau.

### 2.5 Statistics

Statistical analyses are detailed in the Supplementary Methods, and were performed in SPSS 26.0 (Chicago, IL). Cortical thickness between groups and its association with pathology was tested with linear mixed models (LMM) with age, gender and post-mortem delay as covariates, and Bonferroni post-hoc correction for multiple testing. Pathological outcome measures were compared across groups with LMM, using age and gender as covariates. Statistics at the brain area level were corrected for multiple comparisons using the false discovery rate (FDR) approach^30^, and the corrected p-values were expressed as q-values.

## 3. RESULTS

### 3.1 Donor characteristics

Demographical, clinical, radiological, and pathological data of AD and non-neurological control donors are summarized in Table 1. Age, disease duration and post-mortem delay did not differ between groups, whereas gender differed between controls and AD cases (p=0.032). On MRI, normalized brain volume (−5.4% in AD compared to controls, p=0.040), and normalized grey matter volume (−11.6%, p=0.001), but not normalized white matter volume (p=0.735) were lower in AD cases compared to controls. As per definition, AD cases had higher Braak NFT stage (p<0.001), Thal phase (p<0.001) and ABC score (p<0.001) than controls. Regarding AD phenotypes, typical and atypical AD did not differ in any demographical, radiological, and pathological data. APOE genotype did not differ between AD phenotypes (p=0.315).

**Table 1.** Clinical, radiological and pathological characteristics of included donors.

|  | Control | AD | Typical AD | Atypical AD |
| --- | --- | --- | --- | --- |
| **Clinical characteristics** |  |  |  |  |
| **N (phenotype)** | 10 | 19 | 10 | 9 (6 B/D, 3 PCA) |
| **Gender M/F (%M)** | 4/6 (40%) | 16/3 (84%) * | 9/1 (90%) | 7/2 (78%) |
| **APOE genotype N** | 9 | 19 | 10 | 9 |
| ε4 non-carrier | 5 (56%) | 7 (37%) | 5 (50%) | 2 (22%) |
| ε4 heterozygous | 4 (44%) | 10 (53%) | 4 (40%) | 6 (67%) |
| ε4 homozygous | - | 2 (10%) | 1 (10%) | 1 (11%) |
| **Age at disease onset**<br>years, mean ± SD | - | 60 ± 10 | 61 ± 8 | 59 ± 13 |
| **Age at death**<br>years, mean ± SD | 69 ± 7 | 67 ± 12 | 70 ± 11 | 64 ± 12 |
| **Disease duration**<br>years, mean ± SD | - | 8 ± 5 | 10 ± 6 | 6 ± 3 |
| **Post-mortem delay**<br>minutes, mean ± SD | 549 ± 114 | 478 ± 116 | 520 ± 95 | 432 ± 124 |
| **Radiologic characteristics** |  |  |  |  |
| **NBV** (L) mean ± SD | 1.49 ± 0.06 | 1.41 ± 0.13 * | 1.42 ± 0.13 | 1.40 ± 0.14 |
| **NGMV** (L) mean ± SD | 0.76 ± 0.04 | 0.67 ± 0.09 ** | 0.68 ± 0.07 | 0.67 ± 0.11 * |
| **NWMV** (L) mean ± SD | 0.72 ± 0.03 | 0.73 ± 0.08 | 0.73 ± 0.09 | 0.73 ± 0.08 |
| **Pathological characteristics** |  |  |  |  |
| **ABC score**<sup>31</sup> N | 10 | 19 | 10 | 9 |
| A 0/1/2/3 | 3/6/1/0 | 0/0/0/19 *** | 0/0/0/10 *** | 0/0/0/9 *** |
| B 0/1/2/3 | 3/7/0/0 | 0/0/4/15 *** | 0/0/3/7 *** | 0/0/1/8 *** |
| C 0/1/2/3 | 10/0/0/0 | 0/0/4/15 *** | 0/0/3/7 *** | 0/0/1/8 *** |
| **Thal phase**<sup>4</sup> N | 10 | 19 *** | 10 *** | 9 ** |
| 0/1/2/3/4/5 | 3/3/3/1/0/0 | 0/0/0/1/1/17 | 0/0/0/1/0/9 | 0/0/0/0/1/8 |
| **Braak NFT stage**<sup>5</sup> N | 10 | 19 *** | 10 *** | 9 ** |
| 0/1/2/3/4/5/6 | 3/6/1/0/0/0/0 | 0/0/0/0/4/8/7 | 0/0/0/0/3/3/4 | 0/0/0/0/1/5/3 |
| **CAA type** N | 10 | 18 | 10 | 9 |
| Type 1 / type 2 (% type 1) | 0/0 | 15/3 (83%) | 7/2 (78%) | 8/1 (89%) |

Abbreviations: N= sample size; B/D= behavioral / dysexecutive variant; PCA= posterior cortical atrophy; M/F= males/females ratio; SD= standard deviation; L= liter; NBV= normalized brain volume; NGMV= normalized grey matter volume; NWMV= normalized white matter volume; NFT= neurofibrillary tangles; CAA= cerebral amyloid angiopathy. \* $p$ <0.05, \*\* $p$ <0.01, \*\*\* $p$ <0.001 when compared to controls.

### 3.2 MRI cortical atrophy of the temporo-parietal region in AD

AD donors had significantly more cortical atrophy compared to controls (−5.7%, p=0.011, see Table S4), in the inferior (q=0.004), middle (q=0.008) and superior temporal gyrus (q=0.012), entorhinal cortex (q=0.001), fusiform gyrus (q=0.050), inferior parietal gyrus (q=0.020), insular cortex (q=0.043), supramarginal cortex (q=0.020) and precuneus (q=0.028) in the left hemisphere, and in the entorhinal cortex (q=0.025) and inferior temporal gyrus (q=0.042) in the right hemisphere (Figure 2A). Compared to controls, atypical AD cases showed significant global cortical atrophy (−6.2%, p=0.039), while typical AD cases did not (p=0.116) (Figure 2B). No significant difference in atrophy was found between AD phenotypes (p=1.000) (Table S4).

**Figure 2.**
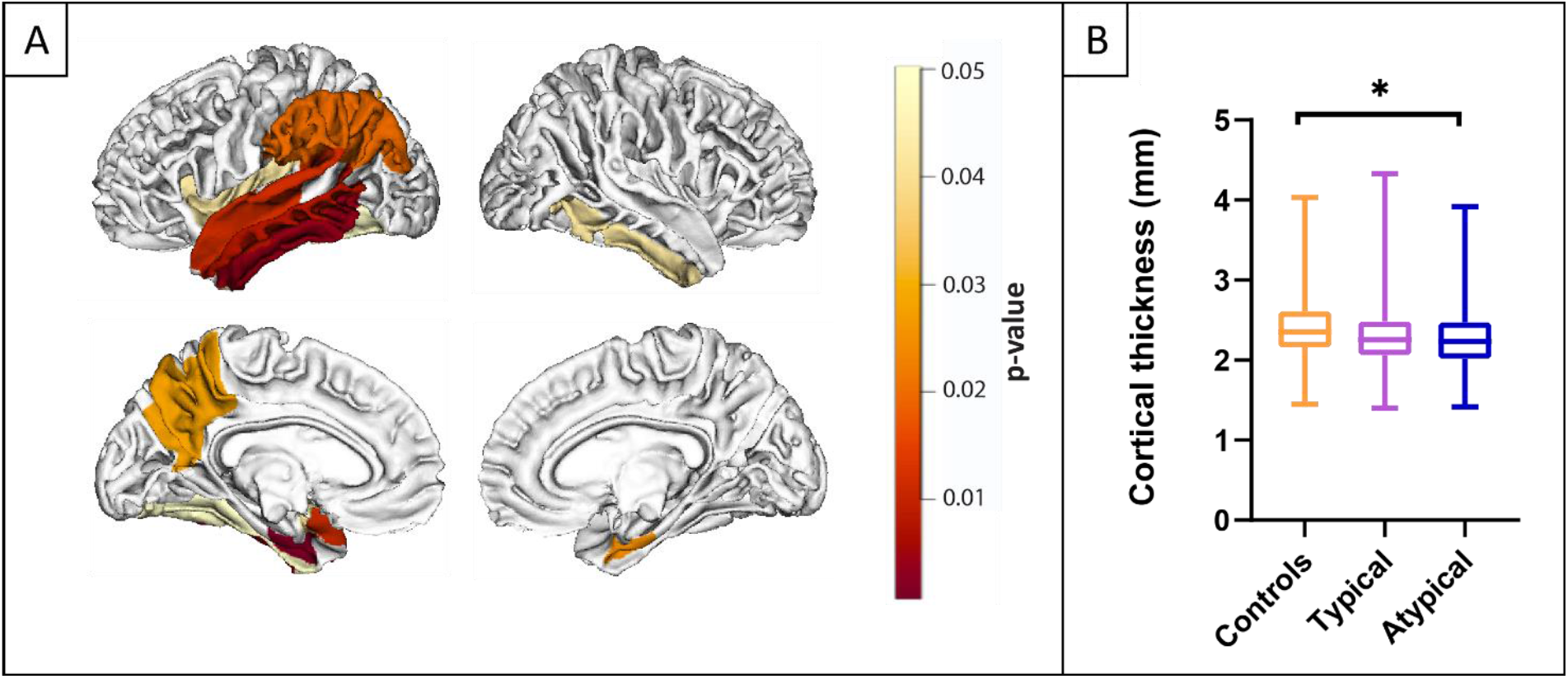
MRI cortical atrophy in AD. Figure (A) shows the atrophic patterns in AD compared to controls (when typical and atypical AD were combined) across the whole cortex. The scale bar represents the p-values. No significant differences in atrophy patterns were found between typical and atypical AD. Graph (B) shows differences in global cortical thickness in controls, typical and atypical AD. The boxplot represents the median, the upper and lower quartile, and the minimum and maximum values. *p<0.05 when compared to controls. For detailed information, see Table S4.

### 3.3 Load and distribution of AD pathological hallmarks

As expected, AD cases had significantly higher Aβ (p<0.001) and p-tau load (p<0.001) compared to controls (see Table S5). By subgroup, both typical and atypical AD had higher Aβ (p=0.012 and p=0.001, respectively) (Figure 3A-C) and p-tau load (p=0.002 and p<0.001, respectively) (Figure 3D-F) compared to controls, whereas typical and atypical AD did not differ in Aβ (p=1.000) or p-tau load (p=1.000).

**Figure 3.**
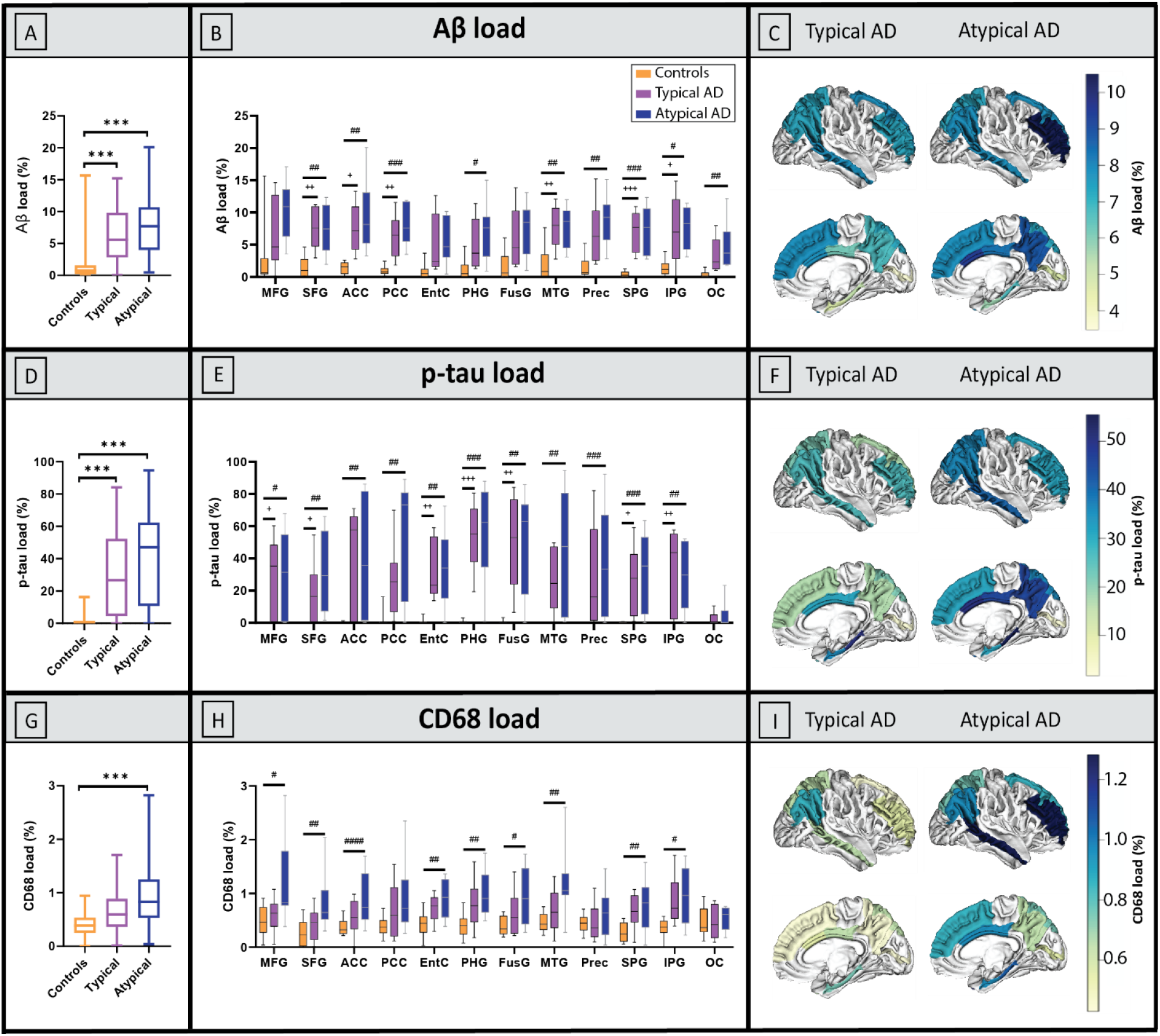
Load and distribution of pathological hallmarks in AD phenotypes and controls. A-C represent the load of Aβ, D-F of p-tau, and G-I of reactive microglia in the right hemisphere. In the first column (A, D and G) show group differences in overall pathological load with boxplots showing median, the upper and lower quartile, and the minimum and maximum values for each group; the middle column (B, E and H) show group differences across regions; C, F and I visually show the mean pathological load on the cortical surface in typical and atypical AD phenotypes, i.e. the same data graphically showed in the middle column. In short, we found significant differences in Aβ and p-tau distribution patterns compared to controls, but not between AD phenotypes. Additionally, atypical AD donors had an overall higher reactive microglia load compared to typical AD, which tended to be higher in all the regions, and the regions with the highest mean reactive microglia values were the frontal and temporal cortex. Legend: MFG: middle frontal gyrus, SFG: superior frontal gyrus, ACC: anterior cingulate cortex, PCC: posterior cingulate cortex, EntC: entorhinal cortex, PHG: parahippocampal gyrus, FusG: fusiform gyrus, MTG: middle temporal gyrus, Prec: precuneus, SPG: superior parietal gyrus, IPG: inferior parietal gyrus, OC: occipital cortex. In the first column: * (p<0.05), ** (p<0.010), *** (p<0.001) when compared to controls. In the middle column, FDR corrected p-values: ^+^ (q<0.05), ^++^ (q<0.010), ^+++^ (q<0.001) typical AD compared to controls; ^#^ (q<0.05), ^###^ (q<0.010), ^###^ (q<0.001) atypical AD compared to controls.

AD cases had a significantly higher reactive microglia load compared to controls (p=0.002, see Table S5). By subgroup, atypical AD cases had a higher reactive microglia load than controls (p=0.001), whereas typical AD cases did not (p=0.104) (Figure 3G-I), suggesting that the significant difference in reactive microglia load between AD and controls was driven by the atypical AD group. No significant difference was found between AD phenotypes (p=0.199). For an overview of the correlations between pathological markers, see table Table S6.

### 3.3 Cortical thickness associates with Thal and Braak staging

In the whole cohort, the average whole-brain cortical thickness correlated negatively with both Thal phase (r_s_ = -0.39, R^2^=15%, p=0.037) and Braak NFT stage (r_s_ = -0.40, R^2^=16%, p=0.030, Figure S1), suggesting an increase in cortical atrophy with disease progression.

### 3.4 Aβ load weakly correlates with less cortical atrophy in AD

A weak positive correlation between Aβ load and cortical thickness was found in the AD group across regions (r = 0.19, R^2^=3%, p=0.010; Table S7, Figure 4A) but not in controls (p=0.165). A similar association was found in typical AD (r = 0.22, R^2^=5%, p=0.022), but not in the atypical AD (p=0.200). We found no associations within brain areas for any group. Overall, Aβ load weakly correlated with less cortical atrophy in AD.

**Figure 4.**
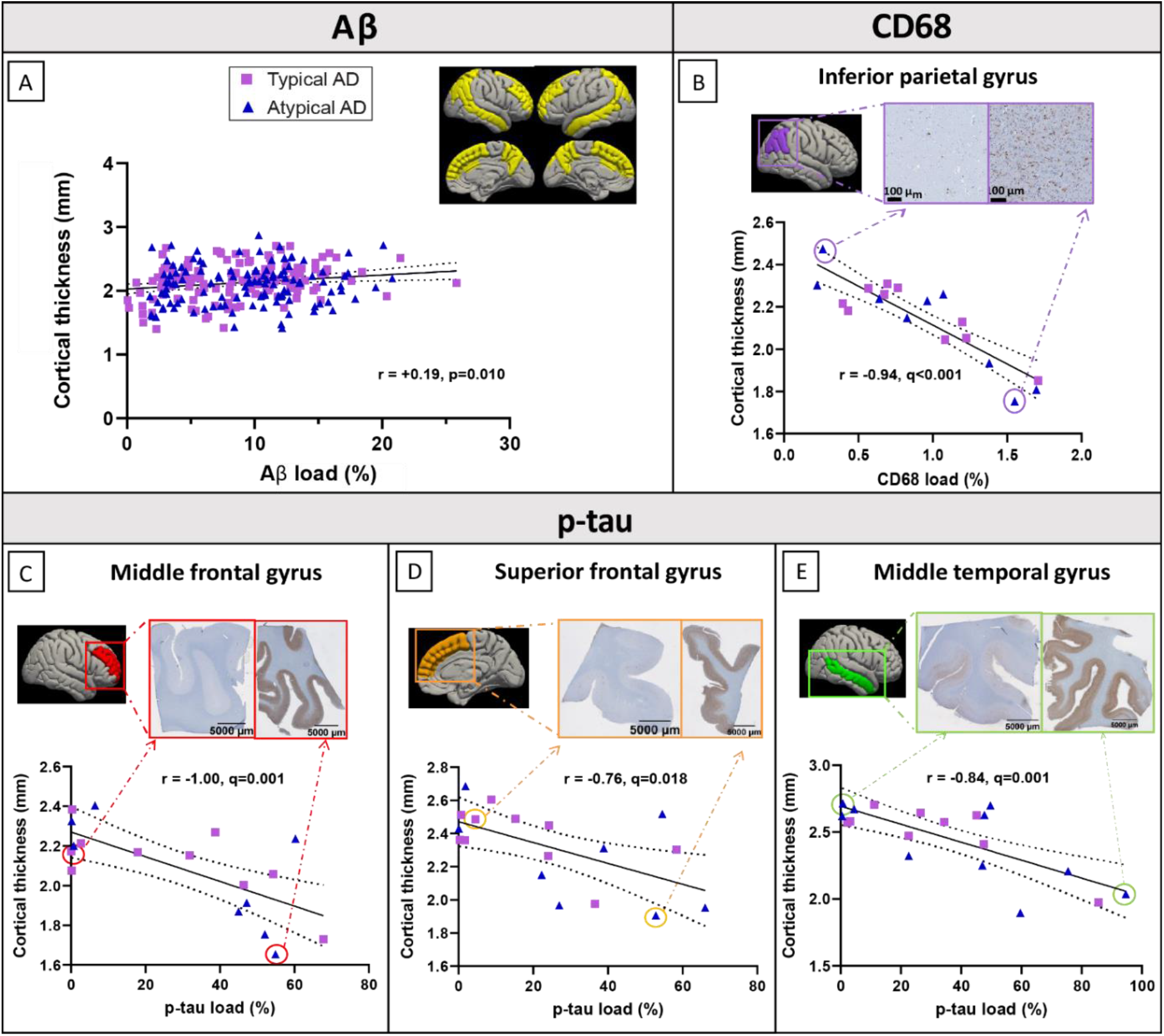
MRI-pathology associations in AD. (A) Weak positive correlation between Aβ load and cortical thickness in the AD group across regions (shown in yellow). (B) Strong negative correlation between reactive microglia load and cortical thickness in the inferior parietal gyrus. (C) Strong negative correlations between p-tau load and cortical thickness in the middle frontal gyrus, (D) superior frontal gyrus, (E) and middle temporal gyrus. On the top right of each graph, two tissue sections representing a low (left) and high (right) pathological load are shown. Purple squares represent typical AD cases, and blue triangles atypical AD cases. A fit-line with 95% confidence interval (dashed lines) is shown for each correlation.

### 3.5 P-tau load strongly correlates with regional cortical atrophy

No associations were found between p-tau load and cortical thickness in AD (p=0.477), AD phenotypes (p=0.144 for typical, and p=0.856 for atypical), or controls (p=0.755) across regions. However, within regions, we observed strong significant correlations in the middle (r = -1.00, R^2^=100%, q=0.001) and superior frontal gyrus (r = -0.76, R^2^=58%, q=0.018), and the middle temporal gyrus (r = -0.84, R^2^=71%, q=0.001) in the AD group only (Table S8, Figure 4C-E). Overall, p-tau load strongly correlated with cortical atrophy in frontal and temporal regions in AD.

### 3.6 Reactive microglia load contributes to cortical atrophy in the parietal region

No associations were found between reactive microglia load and cortical thickness in the AD group (p=0.487) nor controls (p=0.242) across regions. Similarly, we found no associations in typical (p=0.747) and atypical (p=0.285) AD phenotypes. However, within regions, we found a strong and negative significant association in the right inferior parietal gyrus in AD donors (r = -0.94, R^2^=89%, q<0.001) (Figure 4B), which survived also when p-tau load was included in the model as covariate (r = -0.86, R^2^=74%, q<0.001). Overall, reactive microglia load strongly correlated with cortical atrophy in the inferior parietal gyrus in AD.

### 3.7 Combined contribution of pathological hallmarks to cortical thickness

A regression model was run to investigate the combined contribution of Aβ, p-tau and reactive microglia load on cortical thickness of each brain area, and to investigate which had the strongest association with cortical atrophy. While we found no associations in controls, we found significant associations in AD in the middle frontal gyrus (r = 0.88, R^2^=77%, q=0.031), and the inferior parietal gyrus (r = 0.92, R^2^=84%, q<0.001), explaining up to 84% of the variance in cortical thickness. In AD, p-tau load was the major contributor in the middle frontal gyrus (q=0.007), while reactive microglia was the main contributor in the inferior parietal gyrus (q=0.001). When the AD group was split up in typical and atypical AD, no areas showed significant regression models.

### 3.8 From post-mortem *in-situ* to ante-mortem *in-vivo*

Additionally, we investigated the association between ante-mortem *in-vivo* and post-mortem *in-situ* MRI scans of the same donor. Cortical thickness assessment of scans acquired <1 year before death tended to have a stronger correlation with post-mortem cortical thickness than ante-mortem scans acquired 8-10 years before death (r ranged from 0.98 in ante-mortem scan with a 2-year interval from death to 0.69 in ante-mortem scan with a 10-year interval from death, p<0.001 for all, Table S9). Moreover, we investigated the correlation between p-tau load and both ante-mortem and post-mortem cortical thickness in the 3 brain areas that showed significant correlations in our study (see Paragraph 3.5). Cortical thickness measured from ante-mortem scans acquired shortly before death (<1 year, n=3) showed high concordance with post-mortem cortical thickness and its relationship with histopathology, while cortical thickness measured from ante-mortem scans acquired 8-10 years before death (n=2) showed more discordance with post-mortem cortical thickness, especially in the middle temporal gyrus (Figure S2A-B). In one AD case which had 3 ante-mortem scans in addition to the post-mortem scan, we found a progressively increased pattern of discordance with more years between ante-mortem scan and death, with an mean deviation from post-mortem cortical thickness of 22 μm±17 at 1-month, 24 μm±17 at 2-year, and 107 μm±37 at 3-year intervals (Figure S2C).

## 4 DISCUSSION

Using a combined post-mortem *in-situ* MRI and histopathology approach, we investigated the associations between MRI cortical thickness and Aβ, p-tau and reactive microglia load in clinically-defined and pathologically-confirmed AD and control donors, and explored these associations in typical and atypical AD phenotypes. Associations between the histopathological hallmarks and MRI cortical atrophy were found in the AD group, and not in controls. In AD, Aβ and p-tau load contributed differently to cortical thickness: Aβ associated weakly and diffusely with reduced cortical atrophy, while p-tau accumulation strongly associated to regional cortical atrophy. In the cortex of AD donors, the strongest contributors to cortical atrophy were p-tau load in the frontal and temporal cortices, and reactive microglia load in the parietal region.

On MRI, our AD cohort showed pronounced cortical atrophy in the temporoparietal region of the left hemisphere, consistent with the classic AD signature and independent of the phenotypical presentation^2,3,9^.

We found a higher Aβ load in both AD phenotypes compared to controls, but we did not observe a difference in Aβ load nor distribution between AD phenotypes, which is in line with previous studies^8,9^. Furthermore, we found a high p-tau load in AD excluding the occipital cortex, consistent with the fact that this region is the last affected according to Braak NFT staging^5^ and 12 out of 19 of our cases not having reached this stage yet. We did not find any significant differences in p-tau load nor distribution in clinically-defined AD phenotypes, although a difference in distribution has been described previously^8,32–34^. However, most of those studies used pathologically-defined AD donors as opposite to our clinically-defined cohort^8,33,34^, and the only study that used clinically-defined phenotypes did not find any significant difference in NFT load or distribution in the B/D variant compared to typical AD cases^32^, which represents 6 out of 9 cases of our atypical AD cohort.

In line with the literature^6,7^, we show that AD cases had a higher reactive microglia load compared to controls, which was driven by the atypical AD group in our cohort. Atypical AD cases had a higher reactive microglia load compared controls, while typical AD cases did not. Neuroinflammation tends to be stronger in relatively young AD patients compared to the oldest patients^35^, however our atypical cohort did not have a younger age compared to our typical AD group. As such, the underlying disease mechanisms that lead to an increased inflammatory response remains to be elucidated.

When associating our MRI findings with AD pathological hallmarks, we found a weak positive association between Aβ load and cortical thickness in AD, suggesting that a higher Aβ load associated to a slightly decreased global cortical atrophy (Figure 5). Previous studies also reported that Aβ deposition is widespread across the cortex in AD, however they were not consistent in the association with cortical atrophy nor in the direction of the association; some PET studies showed no correlation between Aβ and cortical thickness or grey matter volume^9,11,12,36^, while other cerebrospinal fluid (CSF) studies found a positive correlation^37–41^. The latter argued the existence of a “two-phase phenomenon” along the AD continuum, according to which cortical thickness follows a biphasic trajectory: in preclinical phases there is a cortical thickening, suggesting a relationship with amyloid deposition, and in the late clinical phases cortical atrophy occurs, indicating the (additional) influence of p-tau accumulation^37–42^. Our results being similar to these latter CSF studies is not due to the inclusion of preclinical AD cases, since the positive association was found in our clinical AD group, but more likely due to the use of immunohistochemical quantification. In fact, immunohistochemistry is sensitive to pick up even small diffuse accumulations of Aβ (which is limited with PET imaging^43–45^). Therefore, it is possible that our study revealed a positive association between Aβ and cortical thickness in late clinical AD similar to preclinical AD.

**Figure 5.**
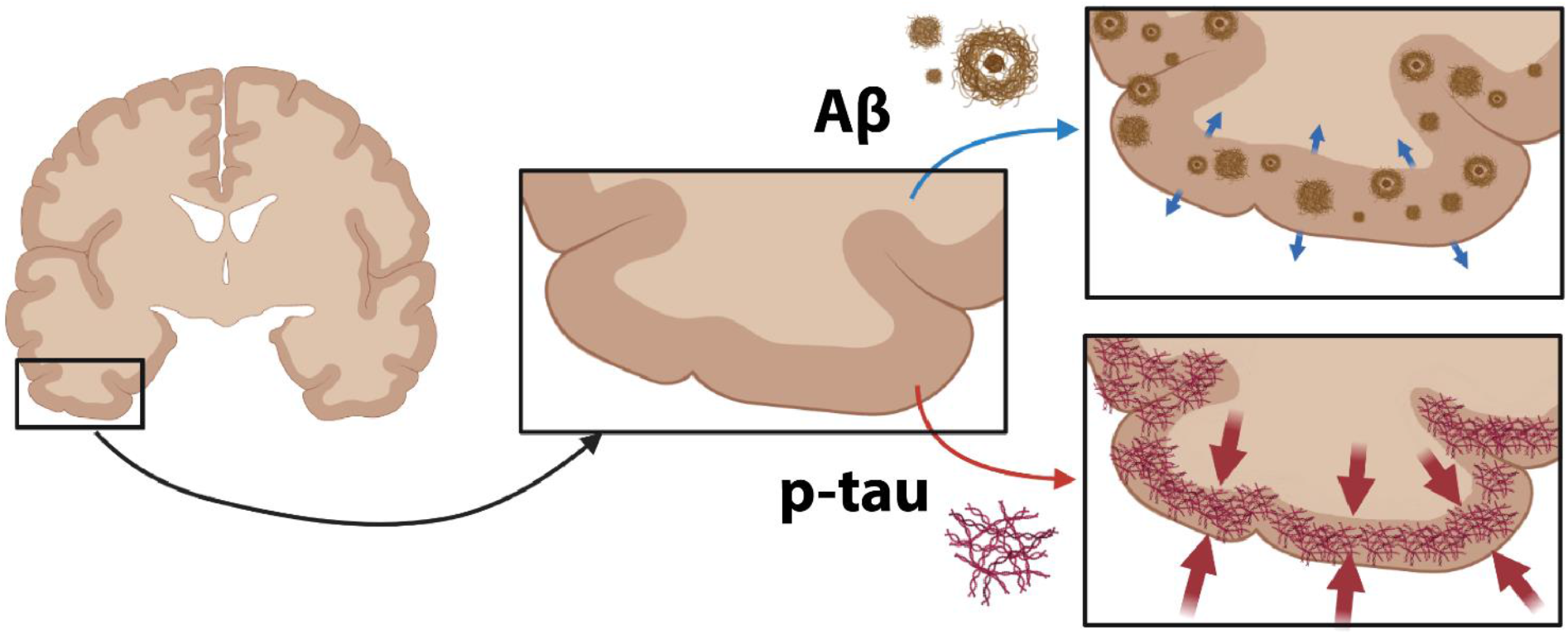
Summary figure: differential effects of p-tau and Aβ load on cortical thickness in AD. Aβ weakly correlates with a reduced cortical atrophy (top; small blue arrows), while p-tau load strongly correlates with cortical atrophy (bottom; big red arrows) in AD.

We found that the regional increase in p-tau load strongly associated with cortical atrophy in frontal and temporal regions in AD (Figure 5). These results are in line with several PET studies which reported regional correlations between tau PET and cortical thickness or grey matter volume^9–13,36^. Even if the exact mechanisms of p-tau-mediated neurodegeneration are still unclear, p-tau has been shown to be closely related to axonal transports deficits and neuronal and synaptic loss, leading to volume loss, hence cortical atrophy^46^. A striking finding in our study is the variance in cortical thickness explained by p-tau load, which ranges between 58% in the superior frontal gyrus to 100% in the middle frontal gyrus, suggesting that p-tau load is indeed one of the main contributors to cortical atrophy in AD. While temporal regions are expected to be hit by p-tau-associated atrophy in AD^3^, our results indicate that frontal regions are also particularly vulnerable to p-tau pathology, which might be due to our inclusion of atypical AD cases of the B/D phenotype.

Regarding the association between cortical thickness and neuroinflammation, we found a strong association between reactive microglia load and cortical atrophy in the parietal region in AD. Similar findings have been reported in a PET study, where the PET marker ^11^CPK11195 for microglial activation correlated with parieto-occipital thinning in AD, including the inferior parietal gyrus^16^. The temporoparietal region reveals neuroinflammation in AD^47^, which can contribute to structural damage. Since the temporal region is one of the first to be affected^5^, any correlational analysis with structural imaging is likely to suffer from a floor effect in end-stage AD cases^16^. Therefore cortical thinning is, most likely, significantly correlated with neuroinflammation only in parietal areas. When chronically activated, microglia tend to transform to a dystrophic, senescent phenotype, which brings them to lose their neuroprotective functions and to become detrimental and neurotoxic, thus accelerating the disease course^7^. Senescent reactive microglia are known to release cytokines, reactive oxygen species and pro-inflammatory factors^6^, which can contribute to synaptic and neuronal loss^7^, hence neurodegeneration.

To further validate our results, we explored the correlation between post-mortem and ante-mortem cortical thickness extrapolated from scans acquired shortly before death and several years prior to it, and its correlation to histopathology. Our findings show that post-mortem *in-situ* scans showed high concordance with ante-mortem *in-vivo* scans acquired few months prior to death. On the other hand, when the ante-mortem scans were acquired several years before death, the associations showed discrepancies. To conclude, this confirms that post-mortem MRI can be used as a proxy for *in-vivo* MRI^48^.

The main strength of this study is that MRI and gold-standard immunohistological data were collected from the same donor at the same moment in time. All donors had pathological confirmation of clinical diagnosis, as clinical-pathological discrepancies occur in 10% of cases^49^, and may obscure *in-vivo* studies. In addition, comprehensive clinical and pathological datasets were available, and both ante-mortem and post-mortem MRI were collected for a subset of patients, making this study encompassing clinical, pathological and radiological data. However, there are also limitations, such as the small group sample sizes, the heterogeneity of our atypical AD cohort, and the fact that the ante-mortem MRI scans have not been collected systematically, having therefore slightly different parameters. Additionally, since we used only one antibody for p-tau, glial and neuronal tau could not be differentiated, and similarly, different Aβ variants were not differentiated. Moreover, we investigated three markers that might contribute to cortical thickness changes, while it is likely that also other cellular and molecular components contribute to cortical atrophy, such as synaptic and axonal degeneration. Future research should therefore investigate the MRI-pathology associations in a larger cohort and include more neurodegenerative markers and molecular profiling to further validate MRI atrophy patterns in AD.

Taken together, our findings show that in AD Aβ load correlates to a reduced global cortical atrophy, while p-tau load is the strongest contributor to regional cortical atrophy in fronto-temporal regions, and reactive microglia load is a strong correlate of cortical atrophy in the parietal region. An exploration within AD phenotypes showed an increased reactive microglia load in atypical AD, but not typical AD, compared to controls, even though no or only subtle differences were found in MRI-pathology associations, indicating that the histopathological correlates of cortical atrophy might be similar between AD phenotypes included in our study. Moreover, we show that post-mortem *in-situ* MRI can be used as proxy for ante-mortem *in-vivo* MRI. In conclusion, our results show that distinct histopathological markers correlate differently with cortical atrophy, highlighting their different roles in the neurodegenerative process, therefore contributing to the understanding of the pathological underpinnings of MRI atrophic patterns.

## Acknowledgments

We would like to thank all brain donors and their next of kin for brain donation, the Netherlands Brain Bank (NBB)and Normal Aging Brain collection Amsterdam (NABCA) autopsy teams, Tjado HJ Morrema and Danae de Gooijer for cutting sections and Martijn Steenwijk for the lesion filling MRI script. FB is supported by the NIHR biomedical research centre at UCLH.

## Conflicts

The authors declare that they have no competing interests.

## Funding sources

This study was funded by PAGE-AD Alzheimer Association (Research Fellowship AARF-18-566,459), ZonMW Memorabel (grant # 733050102), and MJFF (grant # 17253).

## Notes

### Competing Interest Statement

The authors have declared no competing interest.

