## Supplementary material for "Amyloid-β, p-tau, and reactive microglia load are correlates of MRI cortical atrophy in Alzheimer’s disease": Supplemetary figures

### Supplementary figures

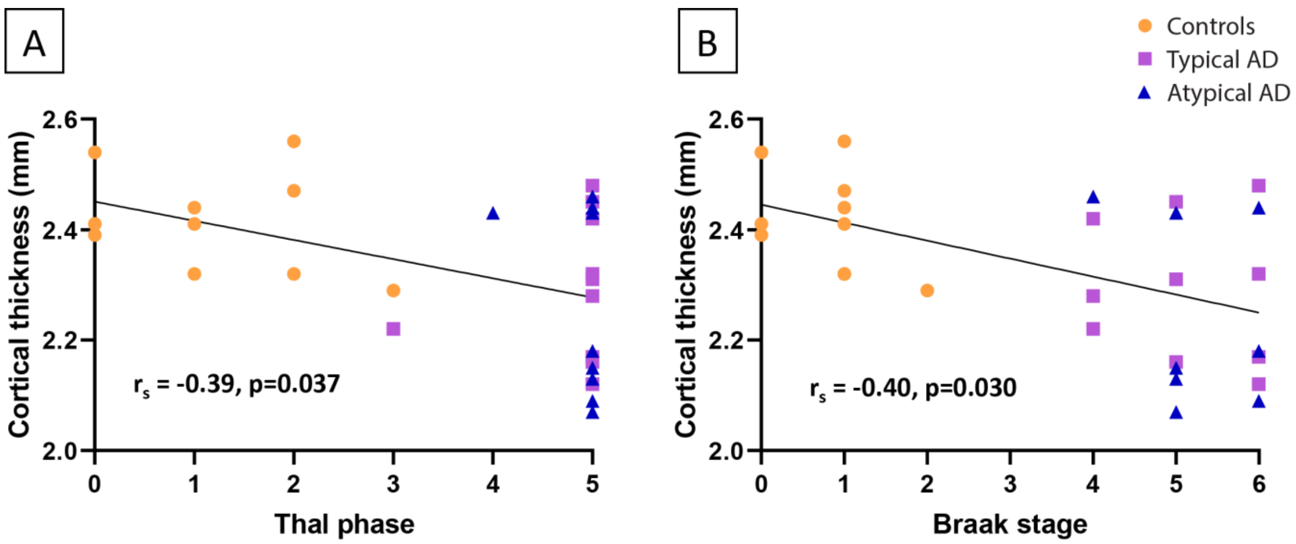


**Figure S1. Association of MRI cortical thickness with Thal phase (A) and Braak NFT stage (B) at autopsy in AD.** Averaged whole-brain cortical atrophy associated with a higher Thal phase[4] and Braak NFT stage[5] in the whole cohort, including controls, typical and atypical AD.

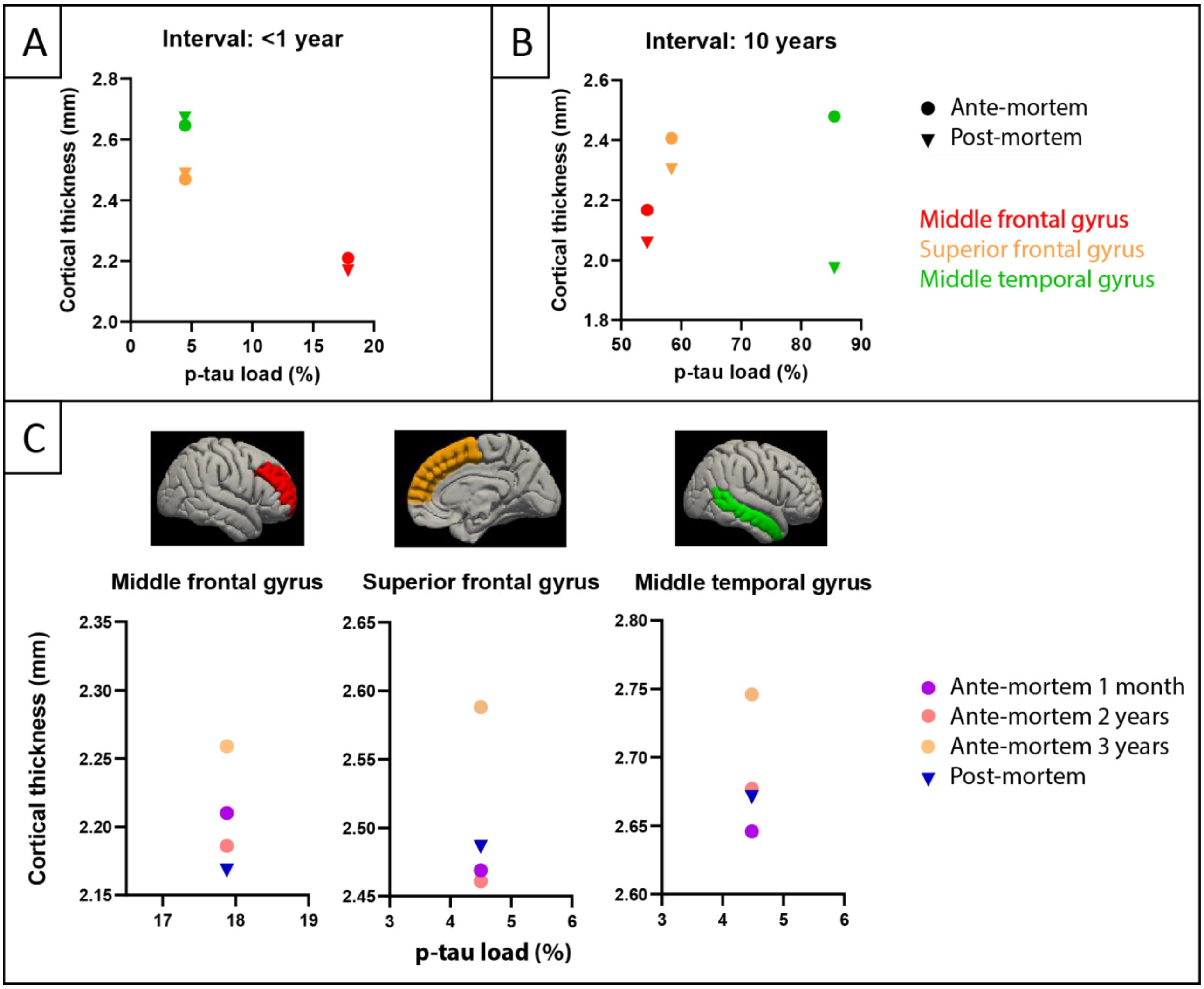


**Figure S2. Association of post-mortem and ante-mortem MRI cortical thickness with p-tau load.** (A) and (B) show the association of p-tau load and both ante-mortem and post-mortem cortical thickness in the regions that hosted a significant association (see Paragraph 3.5). Particularly, (A) shows that the association between ante-mortem and post-mortem cortical thickness with p-tau load does not differ when the ante-mortem scan is acquired less than one year before death (case number 20 in Table S9), (B) while some variation is found when the scan is acquired 10 years before death, particularly in the middle temporal gyrus (case number 17 in Table S9). (C) shows a single AD case (case number 20 in Table S9) with three ante-mortem scans acquired at one-month (purple circle), two-years (pink circle), and three-years intervals (orange circle) prior to death, compared to the post-mortem scan (blue triangle). In each brain area, the three-year ante-mortem scan showed the most pronounced difference in cortical thickness compared to the post-mortem scan.
