## Supplementary methods for "Amyloid-β, p-tau, and reactive microglia load are correlates of MRI cortical atrophy in Alzheimer’s disease"

### 1. MRI acquisition

The following post-mortem sequences were acquired for all subjects: i) a sagittal 3D T1-weighted fast spoiled gradient echo sequence (repetition time (TR) = 7 ms, echo time (TE) = 3 ms, flip angle = 15°, 1-mm-thick axial slices, in-plane resolution= 1.0 x 1.0 $mm^2$); ii) and a sagittal 3D fluid attenuation inversion recovery (FLAIR) sequence (TR = 8000 ms, TE = 130 ms, inversion time (TI) = 2000-2500 ms, 1.2-mm-thick axial slices, in-plane resolution= 1.11 x 1.11 $mm^2$), with TI corrected for post-mortem delay. Subsequently to MRI acquisition, the autopsy was immediately performed, resulting in a total post-mortem delay within 10 hours for all brain donors. Ante-mortem sequences that were acquired for diagnostics were retrospectively obtained from the Amsterdam Dementia Cohort[1], from two different scanners (3T GE MR750 and Philips 3T Achieva). The parameters were slightly different between scanners, with TR varying between 7.8 and 7.9 ms, and TE between 2.9 and 5.2 ms, while voxel size was fixed at $1mm^3$ (**Table S2** for the sequence details).

### 2. Filling of white matter hyperintensities on MRI

Post-mortem T1w images were lesion filled to reduce lesion effects on subsequent automated segmentations. Segmentation of white matter abnormalities was performed on FLAIR images using multi-view convolutional neural network with batch normalization followed by manual editing, obtaining lesion maps, which were registered to the 3D T1 images. The refilling of the lesions was done using LEAP[2].

### 3. MRI cortical thickness assessment

Images underwent inhomogeneity correction, removal of non-brain tissue, and segmentation into grey and white matter. Parcellation of the brain was done using the Desikan-Killany atlas[3]. Cortical thickness was measured as the distance from the grey/white matter boundary to corresponding pial surface. The reconstructed datasets were visually inspected, and segmentation errors were corrected.

### 4. MRI brain volume assessment

For all donors, post-mortem normalized brain volume, normalized grey, and white matter volumes were measured from the T1w images using Structural Image Evaluation, using Normalisation, of Atrophy (SIENAX) (part of FSL 5.0.9; http://fsl.fmrib.ox.ac.uk/), which estimates brain tissue volume normalized for skull size[4].

### 5. Immunohistochemistry (IHC)

The formalin-fixed paraffin-embedded sections were cut and mounted on superfrost+ glass slides (Thermo Scientific, USA). The sections were blocked for endogenous peroxidase using 0.3% hydrogen peroxide and 0.1% sodium azide in phosphate buffer saline (PBS; pH 7.4). The sections were immersed in 10mM Citrate buffer ph 6.0 and heated to 120°C in an autoclave for antigen retrieval. Primary antibodies were diluted (as indicated in **Table S2**) in normal antibody diluent (ImmunoLogic, Duiven, The Netherlands) and incubated overnight at 4°C. Primary antibodies were detected using EnVision (Dako, Glostrup, Denmark). Afterwards, antibodies were visualized using 3.3′-Diaminobenzidine (DAB, Dako) with Imidazole (50 mg DAB, 350 mg Imidazole and 30 uL of $H_2O_2$ per 100 mL of Tris-HCl 30mM, pH 7.6). In between steps, PBS was used to wash the sections. After counterstaining with haematoxylin, the sections were dehydrated and mounted with Entellan (Merck, Darmstadt, Germany). To visualize the line of Gennari in the occipital cortex and therefore identify the striate area (i.e. primary visual cortex), we further performed a Kluver staining on these sections.

### 6. Statistics

Normality was tested, and subsequently demographics between AD and controls, and between controls, typical and atypical AD, were compared using a parametric or non-parametric tests for continuous data, and Fisher exact test for categorical data. The associations between cortical thickness and Thal phases / Braak stages were calculated with Spearman's correlation. MRI-pathology associations across regions were tested with linear mixed models in the AD and control group separately, and then within AD phenotypes with age, gender and post-mortem delay as covariates. In MRI-pathology associations across regions in AD, we included the same regions from the right and left hemisphere that had both pathology data and MRI cortical thickness (i.e. the middle and superior frontal cortex, anterior cingulate gyrus, middle temporal cortex, inferior and superior parietal gyrus, precuneus and occipital cortex from both left and right hemisphere). Since controls did not have pathological data from the left hemisphere, we included all the regions from the right hemisphere except for the entorhinal cortex, as this region was significantly thicker than other brain areas (entorhinal cortical thickness: 3.29 mm ±0.57, other areas: 2.20 mm ±0.33, $p<0.001$) and would drive the associations. To investigate the independent effect of each pathological marker on cortical thickness, we used general linear models. To investigate the combined effect of all pathological markers on cortical thickness we used linear regression analysis.
