## Supplementary tables for "Amyloid-β, p-tau, and reactive microglia load are correlates of MRI cortical atrophy in Alzheimer’s disease"

### Table S1. Donor characteristics

| Case number | Phenotype | Gender | Age at death (years) | Disease duration (years) | PMD (minutes) | Cause of death | NBV (L) | NGMV (L) | ABC score[32] | Thal phase[4] | Braak NFT stage[5] |
|---|---|---|---|---|---|---|---|---|---|---|---|
| **Controls** | | | | | | | | | | | |
| **1** | / | M | 68 | - | 510 | Euthanasia | 1.61 | 0.85 | A1 B1 C0 | 2 | 1 |
| **2** | / | F | 63 | - | 490 | Euthanasia | 1.53 | 0.75 | A0 B0 C0 | 0 | 0 |
| **3** | / | F | 72 | - | 440 | Hart failure | 1.52 | 0.80 | A0 B0 C0 | 0 | 0 |
| **4** | / | F | 69 | - | 765 | Pulmonary embolism | 1.38 | 0.70 | A1 B1 C0 | 1 | 1 |
| **5** | / | M | 59 | - | 480 | Euthanasia | 1.49 | 0.77 | A1 B1 C0 | 2 | 1 |
| **6** | / | M | 77 | - | 685 | Pneumonia | 1.46 | 0.72 | A1 B1 C0 | 1 | 1 |
| **7** | / | F | 78 | - | 600 | Unknown | 1.52 | 0.78 | A1 B1 C0 | 1 | 1 |
| **8** | / | F | 59 | - | 490 | Euthanasia | 1.46 | 0.78 | A0 B0 C0 | 0 | 0 |
| **9** | / | F | 71 | - | 410 | Lung Carcinoma | 1.48 | 0.77 | A1 B1 C0 | 2 | 1 |
| **10** | / | M | 74 | - | 620 | Euthanasia | 1.43 | 0.72 | A2 B1 C0 | 3 | 2 |
| **Typical AD** | | | | | | | | | | | |
| **11** | Typical | M | 60 | 2 | 515 | Euthanasia | 1.55 | 0.76 | A3 B3 C3 | 5 | 6 |
| **12** | Typical | M | 68 | 6 | 555 | Euthanasia | 1.43 | 0.72 | A3 B3 C3 | 5 | 5 |
| **13** | Typical | M | 69 | 11 | 715 | Pulmonary infection | 1.66 | 0.69 | A3 B3 C3 | 5 | 5 |
| **14** | Typical | M | 84 | 13 | 353 | Euthanasia | 1.43 | 0.72 | A3 B2 C2 | 5 | 4 |
| **15** | Typical | F | 80 | 7 | 425 | Euthanasia | 1.44 | 0.71 | A3 B2 C2 | 5 | 4 |
| **16** | Typical | M | 53 | 5 | 540 | Palliative sedation | 1.26 | 0.60 | A3 B3 C3 | 5 | 6 |
| **17** | Typical | M | 64 | 12 | 475 | Dehydration | 1.25 | 0.57 | A3 B3 C3 | 5 | 6 |
| **18** | Typical | M | 84 | 23 | 515 | Euthanasia | 1.25 | 0.61 | A2 B2 C2 | 3 | 4 |
| **19** | Typical | M | 77 | 9 | 545 | Excess of pills | 1.40 | 0.67 | A3 B3 C3 | 5 | 6 |
| **20** | Typical | M | 65 | 7 | 560 | Euthanasia | 1.46 | 0.77 | A3 B3 C3 | 5 | 5 |

| Atypical AD | | | | | | | | | | | |
|---|---|---|---|---|---|---|---|---|---|---|---|
| **21** | PCA | M | 65 | 7 | 470 | Cardiac arrest | 1.51 | 0.77 | A3 B3 C3 | 4 | 5 |
| **22** | B/D | M | 59 | n.a. | 210 | Euthanasia | 1.58 | 0.80 | A3 B3 C3 | 5 | 5 |
| **23** | B/D | F | 78 | 4 | 450 | Dehydration | 1.36 | 0.66 | A3 B3 C3 | 5 | 5 |
| **24** | PCA | M | 62 | 8 | 495 | Palliative sedation | 1.16 | 0.53 | A3 B3 C3 | 5 | 6 |
| **25** | B/D | M | 37 | 5 | 671 | Euthanasia | 1.51 | 0.73 | A3 B3 C3 | 5 | 6 |
| **26** | PCA | M | 67 | 9 | 395 | Dehydration | 1.25 | 0.58 | A3 B3 C3 | 5 | 6 |
| **27** | B/D | M | 77 | 4 | 420 | Euthanasia | 1.54 | 0.77 | A3 B2 C2 | 5 | 4 |
| **28** | B/D | F | 59 | 3 | 335 | Swallowing disorder | 1.32 | 0.65 | A3 B3 C3 | 5 | 5 |
| **29** | B/D | M | 73 | 10 | 440 | Cachexia | 1.37 | 0.49 | A3 B3 C3 | 5 | 5 |

Abbreviations: AD = Alzheimer's disease; PCA = posterior cortical atrophy; B/D = behavioral / dysexecutive variant; M = male; F = female; PMD = post-mortem delay; NBV = normalized brain volume; L = liter; NGMV = normalized grey matter volume.

**Table S2. Information on in-vivo MRI scans.**

| Case number | Scanner | Dimensions | In-plane resolution | TE (ms) | TR (ms) | Interval (years) |
|---|---|---|---|---|---|---|
| **12** | 3T GE MR750 | 176x256x256 | $1mm^3$ | 3 | 7.8 | 2 |
| **12** | 3T GE MR750 | 176x256x256 | $1mm^3$ | 3 | 7.8 | 5 |
| **14** | unknown | 176x256x256 | $1mm^3$ | 3.4 | unknown | 2 |
| **14** | 3T GE MR750 | 176x256x256 | $1mm^3$ | 3 | 7.8 | 7 |
| **16** | 3T GE MR750 | 176x256x256 | $1mm^3$ | 3.2 | 7.8 | 4 |
| **17** | unknown | 192x150x256 | $1mm^3$ | 5.2 | unknown | 10 |
| **18** | 3T GE MR750 | 288x288x180 | $1mm^3$ | 3 | 7.8 | 6 |
| **18** | 3T GE MR750 | 176x256x256 | $1mm^3$ | 3 | 7.8 | 7 |
| **19** | Philips 3T Achieva | 256x256x192 | $1mm^3$ | 4.5 | 7.9 | 2 |
| **19** | Philips 3T Achieva | 256x256x192 | $1mm^3$ | 4.5 | 7.9 | 4 |

| | | | | | | |
|---|---|---|---|---|---|---|
| **20** | Philips 3T Achieva | 256x256x192 | 1mm$^3$ | 4.5 | 7.9 | <1 |
| **20** | Philips 3T Achieva | 256x256x192 | 1mm$^3$ | 4.5 | 7.9 | 2 |
| **20** | 3T GE MR750 | 176x256x256 | 1mm$^3$ | 3.2 | 7.8 | 3 |
| **21** | Philips 3T Achieva | 256x256x192 | 1mm$^3$ | 4.5 | 7.9 | 1 |
| **24** | 3T GE MR750 | 176x256x256 | 1mm$^3$ | 3 | 7.8 | 5 |
| **25** | 3T GE MR750 | 174x512x512 | 1mm$^3$ | 2.9 | 7.8 | <1 |
| **26** | 3T GE MR750 | 176x256x256 | 1mm$^3$ | 3.2 | 7.8 | 5 |
| **27** | Philips 3T Achieva | 256x256x192 | 1mm$^3$ | 4.5 | 7.9 | <1 |
| **28** | 3T GE MR750 | 176x256x256 | 1mm$^3$ | 3.2 | 7.8 | 2 |
| **29** | 3T GE MR750 | 176x256x256 | 1mm$^3$ | 3.2 | 7.8 | 4 |

The table shows the information about the ante-mortem *in-vivo* 3T MRI of 14 out of 19 AD cases included in our post-mortem cohort. In addition, 5 out of 14 cases had more than one ante-mortem MRI scan at different intervals from death, measured in years. Case numbers match those in **Table S1**. Legend: TE = time to echo, TR = repetition time; ms = milliseconds.

**Table S3. Information on primary antibodies.**

| Antibody | Antigen | Species | Origin details | Dilution | Incubation time | Antigen retrieval | Detection method |
|---|---|---|---|---|---|---|---|
| **Aβ, clone 4G8** | Aβ amino acid sequence 17-24 | Mouse igG2b | BioLegend, San Diego, USA | 1:8000 | 4°C overnight | Autoclave Citrate buffer (pH 6.0, 10 minutes) | EnVision |
| **p-tau, clone AT8** | Tau phosphorylated at Ser202 and Thr205 | Mouse igG1 | ThermoFisher, Pittsburgh, USA | 1:800 | 4°C overnight | Autoclave Citrate buffer (pH 6.0, 10 minutes) | EnVision |
| **CD68, clone KP1** | CD68 | Mouse igG1 | Dako, Glostrup, Denmark | 1:1200 | 4°C overnight | Autoclave Citrate buffer (pH 6.0, 10 minutes) | EnVision |

**Table S4. Mean and standard deviation of MRI cortical thickness per brain area in each group**.

| Region | | Controls (n=10) | AD (n=19) | Typical AD (n=10) | Atypical AD (n=9) | Q-value |
|---|---|---|---|---|---|---|
| ctx-lh-bankssts | 1 | 2,26±0,09 | 2,07±0,23 | 2,06±0,24 | 2,08±0,24 | ns |
| ctx-lh-caudalanteriorcingulate | 2 | 2,33±0,24 | 2,31±0,32 | 2,20±0,28 | 2,42±0,33 | ns |
| ctx-lh-caudalmiddlefrontal | 3 | 2,34±0,11 | 2,22±0,22 | 2,22±0,17 | 2,22±0,27 | ns |
| ctx-lh-cuneus | 4 | 1,88±0,11 | 1,86±0,15 | 1,84±0,11 | 1,87±0,18 | ns |
| ctx-lh-entorhinal | 5 | 3,48±0,22 | 2,81±0,50 ** | 2,75±0,53 | 2,87±0,49 | q=0.001 |
| ctx-lh-fusiform | 6 | 2,63±0,13 | 2,40±0,22 * | 2,43±0,24 | 2,37±0,22 | q=0.050 |
| ctx-lh-inferiorparietal | 7 | 2,30±0,13 | 2,11±0,15 * | 2,08±0,17 | 2,13±0,12 | q=0.020 |
| ctx-lh-inferiortemporal | 8 | 2,74±0,14 | 2,50±0,19 ** | 2,49±0,23 | 2,50±0,16 | q=0.004 |
| ctx-lh-isthmuscingulate | 9 | 2,12±0,15 | 2,00±0,18 | 1,96±0,18 | 2,06±0,17 | ns |
| ctx-lh-lateraloccipital | 10 | 2,09±0,11 | 2,07±0,14 | 2,04±0,12 | 2,10±0,15 | ns |
| ctx-lh-lateralorbitofrontal | 11 | 2,74±0,19 | 2,61±0,23 | 2,59±0,20 | 2,62±0,27 | ns |
| ctx-lh-lingual | 12 | 1,98±0,05 | 1,93±0,18 | 1,91±0,17 | 1,96±0,20 | ns |
| ctx-lh-medialorbitofrontal | 13 | 2,37±0,13 | 2,24±0,20 | 2,24±0,19 | 2,24±0,22 | ns |
| ctx-lh-middletemporal | 14 | 2,72±0,12 | 2,45±0,20 ** | 2,42±0,19 | 2,48±0,21 | q=0.008 |
| ctx-lh-parahippocampal | 15 | 2,63±0,22 | 2,41±0,34 | 2,33±0,26 | 2,50±0,42 | ns |
| ctx-lh-paracentral | 16 | 2,21±0,07 | 2,23±0,15 | 2,22±0,14 | 2,24±0,17 | ns |
| ctx-lh-parsopercularis | 17 | 2,36±0,08 | 2,27±0,14 | 2,26±0,15 | 2,29±0,12 | ns |
| ctx-lh-parsorbitalis | 18 | 2,72±0,19 | 2,57±0,24 | 2,52±0,25 | 2,62±0,24 | ns |
| ctx-lh-parstriangularis | 19 | 2,32±0,10 | 2,30±0,19 | 2,26±0,21 | 2,34±0,17 | ns |
| ctx-lh-pericalcarine | 20 | 1,74±0,16 | 1,68±0,15 | 1,67±0,17 | 1,68±0,14 | ns |
| ctx-lh-postcentral | 21 | 2,00±0,11 | 1,97±0,10 | 1,99±0,08 | 1,96±0,11 | ns |
| ctx-lh-posteriorcingulate | 22 | 2,18±0,27 | 2,12±0,22 | 2,13±0,24 | 2,11±0,21 | ns |
| ctx-lh-precentral | 23 | 2,33±0,08 | 2,31±0,16 | 2,31±0,13 | 2,30±0,19 | ns |
| ctx-lh-precuneus | 24 | 2,25±0,14 | 2,07±0,14 * | 2,06±0,16 | 2,08±0,12 | q=0.028 |
| ctx-lh-rostralanteriorcingulate | 25 | 2,62±0,20 | 2,46±0,29 | 2,55±0,29 | 2,36±0,28 | ns |
| ctx-lh-rostralmiddlefrontal | 26 | 2,28±0,13 | 2,12±0,18 | 2,12±0,17 | 2,11±0,21 | ns |
| ctx-lh-superiorfrontal | 27 | 2,44±0,13 | 2,33±0,22 | 2,33±0,19 | 2,32±0,26 | ns |
| ctx-lh-superiorparietal | 28 | 2,09±0,12 | 1,96±0,12 | 1,97±0,10 | 1,94±0,14 | ns |
| ctx-lh-superiortemporal | 29 | 2,53±0,14 | 2,37±0,17 * | 2,37±0,17 | 2,37±0,18 | q=0.012 |
| ctx-lh-supramarginal | 30 | 2,34±0,14 | 2,13±0,15 * | 2,15±0,16 | 2,12±0,15 | q=0.020 |
| ctx-lh-frontalpole | 31 | 2,63±0,19 | 2,47±0,27 | 2,42±0,29 | 2,52±0,25 | ns |

| | | | | | | |
|---|---|---|---|---|---|---|
| ctx-lh-temporalpole | 32 | 3,64±0,22 | 3,24±0,41 | 3,24±0,47 | 3,25±0,37 | ns |
| ctx-lh-transversetemporal | 33 | 2,11±0,12 | 2,12±0,21 | 2,10±0,16 | 2,14±0,25 | ns |
| ctx-lh-insula | 34 | 2,71±0,08 | 2,54±0,21 * | 2,60±0,22 | 2,47±0,19 | q=0.043 |
| ctx-rh-bankssts | 35 | 2,30±0,10 | 2,16±0,23 | 2,19±0,24 | 2,12±0,23 | ns |
| ctx-rh-caudalanteriorcingulate | 36 | 2,25±0,22 | 2,22±0,24 | 2,2±0,186 | 2,18±0,31 | ns |
| ctx-rh-caudalmiddlefrontal | 37 | 2,34±0,15 | 2,19±0,20 | 2,20±0,17 | 2,18±0,25 | ns |
| ctx-rh-cuneus | 38 | 1,86±0,08 | 1,88±0,17 | 1,88±0,20 | 1,87±0,13 | ns |
| ctx-rh-entorhinal | 39 | 3,67±0,25 | 3,09±0,60 * | 3,10±0,73 | 3,08±0,45 | q=0.025 |
| ctx-rh-fusiform | 40 | 2,66±0,15 | 2,45±0,26 | 2,48±0,25 | 2,41±0,29 | ns |
| ctx-rh-inferiorparietal | 41 | 2,31±0,11 | 2,15±0,19 | 2,16±0,15 | 2,13±0,24 | ns |
| ctx-rh-inferiortemporal | 42 | 2,79±0,15 | 2,54±0,24 * | 2,63±0,18 | 2,44±0,27 | q=0.042 |
| ctx-rh-isthmuscingulate | 43 | 2,10±0,24 | 2,03±0,24 | 2,11±0,19 | 1,94±0,27 | ns |
| ctx-rh-lateraloccipital | 44 | 2,15±0,08 | 2,14±0,16 | 2,15±0,15 | 2,13±0,17 | ns |
| ctx-rh-lateralorbitofrontal | 45 | 2,72±0,17 | 2,61±0,23 | 2,70±0,16 | 2,51±0,26 | ns |
| ctx-rh-lingual | 46 | 1,98±0,11 | 1,95±0,15 | 1,93±0,16 | 1,98±0,15 | ns |
| ctx-rh-medialorbitofrontal | 47 | 2,48±0,1 | 2,29±0,25 | 2,34±0,24 | 2,23±0,27 | ns |
| ctx-rh-middletemporal | 48 | 2,70±0,08 | 2,45±0,26 | 2,52±0,21 | 2,37±0,30 | ns |
| ctx-rh-parahippocampal | 49 | 2,69±0,35 | 2,40±0,36 | 2,34±0,39 | 2,46±0,33 | ns |
| ctx-rh-paracentral | 50 | 2,23±0,12 | 2,28±0,12 | 2,28±0,15 | 2,29±0,09 | ns |
| ctx-rh-parsopercularis | 51 | 2,32±0,14 | 2,24±0,20 | 2,29±0,15 | 2,19±0,24 | ns |
| ctx-rh-parsorbitalis | 52 | 2,63±0,11 | 2,51±0,24 | 2,55±0,21 | 2,47±0,27 | ns |
| ctx-rh-parstriangularis | 53 | 2,28±0,07 | 2,21±0,17 | 2,23±0,14 | 2,18±0,21 | ns |
| ctx-rh-pericalcarine | 54 | 1,62±0,13 | 1,67±0,15 | 1,68±0,16 | 1,66±0,14 | ns |
| ctx-rh-postcentral | 55 | 1,96±0,08 | 1,95±0,13 | 1,99±0,13 | 1,90±0,12 | ns |
| ctx-rh-posteriorcingulate | 56 | 2,06±0,33 | 2,10±0,27 | 2,21±0,18 | 1,97±0,30 | ns |
| ctx-rh-precentral | 57 | 2,30±0,11 | 2,30±0,15 | 2,30±0,17 | 2,29±0,15 | ns |
| ctx-rh-precuneus | 58 | 2,28±0,13 | 2,11±0,20 | 2,12±0,17 | 2,10±0,23 | ns |
| ctx-rh-rostralanteriorcingulate | 59 | 2,70±0,20 | 2,58±0,36 | 2,63±0,27 | 2,53±0,45 | ns |
| ctx-rh-rostralmiddlefrontal | 60 | 2,26±0,09 | 2,08±0,22 | 2,12±0,18 | 2,04±0,26 | ns |
| ctx-rh-superiorfrontal | 61 | 2,46±0,13 | 2,33±0,24 | 2,38±0,18 | 2,27±0,29 | ns |
| ctx-rh-superiorparietal | 62 | 2,08±0,13 | 1,97±0,16 | 2,01±0,09 | 1,93±0,20 | ns |
| ctx-rh-superiortemporal | 63 | 2,55±0,10 | 2,38±0,23 | 2,43±0,20 | 2,33±0,27 | ns |
| ctx-rh-supramarginal | 64 | 2,33±0,14 | 2,20±0,18 | 2,27±0,10 | 2,14±0,23 | ns |
| ctx-rh-frontalpole | 65 | 2,63±0,27 | 2,38±0,28 | 2,40±0,27 | 2,36±0,30 | ns |

| | | | | | | |
|---|---|---|---|---|---|---|
| ctx-rh-temporalpole | 66 | 3,80±0,16 | 3,34±0,50 | 3,38±0,51 | 3,29±0,51 | ns |
| ctx-rh-transversetemporal | 67 | 2,18±0,14 | 2,16±0,29 | 2,13±0,11 | 2,20±0,19 | ns |
| ctx-rh-insula | 68 | 2,69±0,10 | 2,54±0,29 | 2,61±0,22 | 2,47±0,34 | ns |
| **Whole cortex** | | **2,42 ± 0.43** | **2,28 ± 0.38*** | **2,29 ± 0.38** | **2,27 ± 0.39*** | |

Data are presented as mean ± standard deviation. Regions selected for the study are highlighted in light grey. Legend: ctx = cortex; FDR = false-discovery rate; lh = left hemisphere; rh = right hemisphere. *q<0.05, **q<0.01, when compared to controls, ns: not significant.

**Table S5. Mean and standard deviation of pathological hallmarks**.

| Pathological marker | Controls | AD | Typical AD | Atypical AD |
|---|---|---|---|---|
| **Aβ load (%)** | 1.32 ± 2.04 | 7.03 ± 4.07 *** | 6.45 ± 4.08 * | 7.66 ± 3.99 ** |
| **p-tau load (%)** | 0.36 ± 1.66 | 33.81 ± 27.76 *** | 28.76 ± 26.28 ** | 39.58 ± 28.39 *** |
| **CD68 load (%)** | 0.40 ± 0.21 | 0.77 ± 0.47 ** | 0.64 ± 0.37 | 0.91 ± 0.53 ** |

. Averaged data from brain regions of the right hemisphere. All the values are expressed in mean ± standard deviation. Abbreviations: AD: Alzheimer's disease. *p<0.05, **p<0.01, ***p<0.001 when compared to controls.

**Table S6. Associations between pathological markers in different groups.**

| Pathological marker | Aβ load | p-tau load | CD68 load |
|---|---|---|---|
| **Controls** | | | |
| **p-tau load** | p=0.701 | - | p=0.428 |
| **CD68 load** | r=0.27, p=0.018 | p=0.428 | - |
| **AD** | | | |
| **p-tau load** | p=0.909 | - | r=0.32, p<0.001 |
| **CD68 load** | p=0.408 | r=0.32, p<0.001 | - |
| **Typical AD** | | | |
| **p-tau load** | p=0.660 | - | r=0.18, p=0.029 |
| **CD68 load** | p=0.888 | r=0.18, p=0.029 | - |
| **Atypical AD** | | | |
| **p-tau load** | p=0.767 | - | r=0.40, p<0.001 |
| **CD68 load** | p=0.295 | r=0.40, p<0.001 | - |

The associations between histopathological markers in controls, AD, typical and atypical AD are shown in all the brain areas combined. In each cell, if the correlation was significant, the correlation coefficient (r) of the association and its the p-value are shown.

**Table S7. Correlations between global cortical thickness and Aβ load.**

| Group | r | $R^2$ | P-value |
|---|---|---|---|
| Controls | - | - | p=0.165 |
| AD | 0.19 | 3% | p=0.010 |
| Typical AD | 0.22 | 5% | p=0.022 |
| Atypical AD | - | - | p=0.200 |

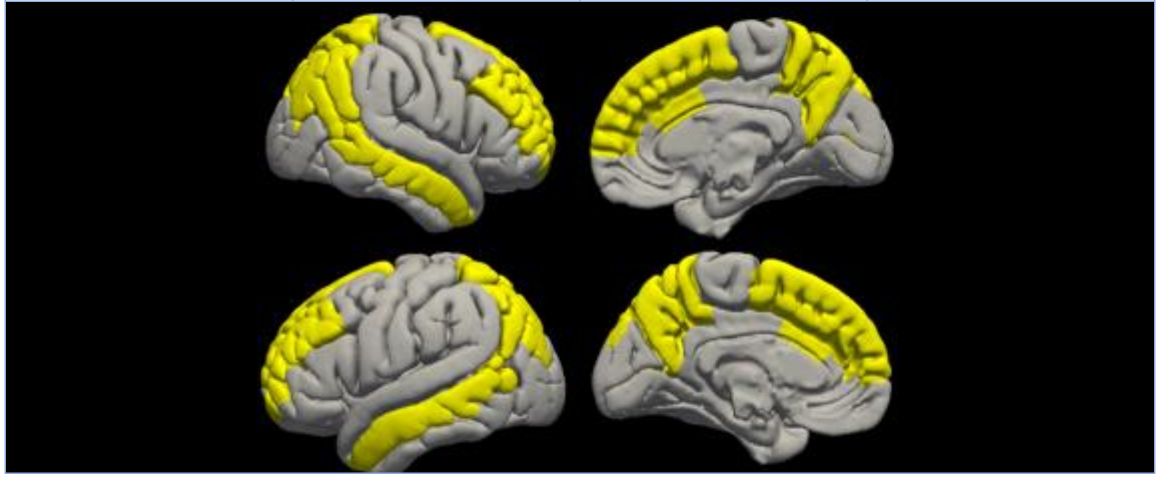

The table shows the associations of cortical thickness with Aβ load in AD, controls, typical AD and atypical AD. The columns show the correlation coefficient (r), explained variance in cortical thickness by Aβ load ($R^2$) and p-value of the correlations tested in the brain regions shown in the last column (in yellow), which were the same for left and right hemisphere: middle frontal and superior frontal gyrus, anterior cingulate cortex, middle temporal gyrus, superior and inferior parietal gyrus, precuneus and occipital cortex.

**Table S8. Correlations between regional cortical thickness on MRI and p-tau load in AD.**

| Region | r | $R^2$ | Q-value | Regional r values |
|---|---|---|---|---|
| Middle frontal gyrus | -1.00 | 100% | q=0.001 | 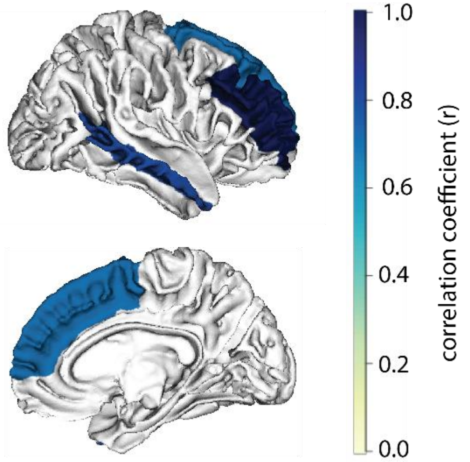 |
| Superior frontal gyrus | -0.76 | 58% | q=0.018 | |
| Anterior cingulate cortex | - | - | ns | |
| Posterior cingulate cortex | - | - | ns | |
| Middle temporal gyrus | -0.84 | 71% | q=0.001 | |
| Entorhinal cortex | - | - | ns | |
| Parahippocampal gyrus | - | - | ns | |
| Fusiform gyrus | - | - | ns | |
| Precuneus | - | - | ns | |
| Superior parietal gyrus | - | - | ns | |
| Inferior parietal gyrus | - | - | ns | |
| Occipital cortex (V1) | - | - | ns | |

The table shows the association of cortical thickness with the p-tau load in regions of the right hemisphere in AD. The columns show the correlation coefficient (r), explained variance in cortical thickness by p-tau load ($R^2$) and q-value (FDR corrected p-value) of the correlations. In the right column, the regional correlation coefficient values (r) indicated in the second column are visually shown on the cortical surface. The table shows that the middle and superior frontal gyrus and the middle temporal gyrus hosted strong, significant negative associations between p-tau load and cortical thickness. Abbreviations: ns = not significant.

**Table S9. Correlations between post-mortem *in-situ* and ante-mortem *in-vivo* cortical thickness in AD cases.**

| Case number | Phenotype | Interval (years) | Interval (months) | r | p-value |
|---|---|---|---|---|---|
| **25** | Atypical (B/D) | <1 | <1 | 0.95 | p<0.001 |
| **20** | Typical | <1 | 1 | 0.97 | p<0.001 |
| **27** | Atypical (B/D) | <1 | 3 | 0.97 | p<0.001 |
| **21** | Atypical (PCA) | 1 | 17 | 0.95 | p<0.001 |
| **20** | Typical | 2 | 22 | 0.98 | p<0.001 |
| **28** | Atypical (B/D) | 2 | 24 | 0.89 | p<0.001 |
| **12** | Typical | 2 | 26 | 0.96 | p<0.001 |
| **14** | Typical | 2 | 28 | 0.96 | p<0.001 |
| **19** | Typical | 2 | 28 | 0.86 | p<0.001 |
| **20** | Typical | 3 | 30 | 0.94 | p<0.001 |
| **16** | Typical | 4 | 48 | 0.91 | p<0.001 |
| **29** | Atypical (B/D) | 4 | 50 | 0.87 | p<0.001 |
| **19** | Typical | 4 | 52 | 0.87 | p<0.001 |
| **26** | Atypical (PCA) | 5 | 54 | 0.95 | p<0.001 |
| **12** | Typical | 5 | 60 | 0.88 | p<0.001 |
| **24** | Atypical (PCA) | 5 | 63 | 0.73 | p<0.001 |
| **18** | Typical | 6 | 72 | 0.94 | p<0.001 |
| **14** | Typical | 7 | 82 | 0.92 | p<0.001 |
| **18** | Typical | 7 | 89 | 0.90 | p<0.001 |
| **17** | Typical | 10 | 120 | 0.69 | p=0.001 |

The table shows the correlations between cortical thickness acquired post-mortem (and used for this study) and ante-mortem at different time intervals from death, measured both in months and years. Case numbers match those in Table S1. Legend: B/D = behavioral / dysexecutive variant; PCA = posterior cortical atrophy; r = correlation coefficient.
